# A failure to replicate the effect of visual symmetry on subjective duration

**DOI:** 10.1101/2020.08.20.258871

**Authors:** Alexis D.J. Makin, Afzal Rahman, Marco Bertamini

## Abstract

Previous work has shown that symmetrical or regular stimuli are judged as lasting longer than asymmetrical or irregular ones, even when actual duration is matched. This effect has been replicated with different methods and stimuli types, both in the UK and Japan. We aimed to a) replicate the effect of symmetry on subjective duration, and b) assess whether it was further modulated by the number of symmetrical axes. There was no evidence for either effect. This null result cannot be explained by reduced statistical power or enhanced floor or ceiling effects. There is no obvious stimulus-based explanation either. However, we are mindful of the reproducibility crisis and file drawer problems in psychology. Other symmetry and time perception researchers should be aware of this null result.

## Introduction

### The scientific value of null results

Many leading researchers doubt general trustworthiness of peer-reviewed science (Bishop, 2019; Ioannidis, 2005). Psychology faces acute criticism because scientific malpractice is allegedly routine, and because there have been famous attempts to estimate reproducibility. Approximately just 39% of published results in psychology are reproducible, and this only increases 51% in cognitive psychology (Open Science Collaboration, 2015). One factor contributing to the replication crisis is well understood: Scientists publish or perish, and the only readily publishable narrative has the standard happy ending: ‘the expected effect was statistically significant (p < 0.05)’. Consequently, the proverbial file drawer accumulates null results, and the literature accumulates false positives (alongside the real effects). Even without blatant data fraud, p-hacking or HARKing (*Hypothesising After Knowing Results*), the published record is likely biased towards ‘lucky’ experiments that overestimate true effect size (Brysbaert, 2019). Such systemic problems have been obvious for decades, but conveniently ignored.

Keith Laws (2013) provides but one useful and provocative commentary. He quotes Francis (2012):

> *“The scientific method is supposed to be able to reveal truths about the world, and the reliability of empirical findings is supposed to be the final arbiter of science; but this method does not seem to work in experimental psychology as it is currently practiced.”*
>
> — *(page 3)*

While this particular comment falls on pessimistic end of the spectrum, even the most optimistic commentators acknowledge substantial room for improvement. Laws (2013) encourages practicing scientists to show leadership and embrace the challenges. However:

> *“This leadership will however require psychologists to take a more active role in submitting replications and null findings – science is clearly not self-correcting.”*
>
> — *(page 7)*

Laws (2013) further recommends that ‘failed’ experiments should not be marginalized in specialist null results journals:

> *“Although laudable, such journals create a special space for replications and null findings rather than acknowledging their place in the centre of science.”*
>
> — *(page 7)*

Following this advice, the current article reports our failure to replicate an effect that published in three different papers, including two from our research group, and one from a Japanese team (Ogden, Makin, Palumbo, & Bertamini, 2016; Palumbo, Ogden, Makin, & Bertamini, 2015; Sasaki & Yamada, 2017).

### The published effect: Symmetry reportedly elongates subjective duration

Ogden et al. (2016) reported that symmetrical patterns were misperceived as staying on the screen longer than random patterns. In their Experiment 1, participants were presented with checkerboard patterns for durations of 500, 750, 1000, 1250 or 1500 ms. The 10 × 10 checkerboards had 40 black and 60 white squares. The arrangements were either 1) two-fold reflectional symmetry, 2) 90-degree rotational symmetry, or 3) random. Participants estimated duration by entering a duration estimate with a computer keyboard. Reflection was judged to have lasted longer than rotation or random (F (2,42) = 17.114, p < 0.001, η_p_^2^ = 0.451). This was replicated in Experiment 2 with an alternative temporal bisection procedure and non-parametric statistical tests.

In an earlier study with a more general research question, Palumbo et al. (2015) also found that symmetrical patterns were judged longer than random patterns. Furthermore, in another study from Waseda University in Tokyo, Sakaki and Yamada (2017) found that high regularity grids were judged as lasting longer than medium or low regularity grids. This is conceptually similar to the effects reported by Palumbo et al. (2015) and Ogden et al. (2016).

Given this consistent literature, we were confident that the visual symmetry lengthens subjective duration. The effect could be mediated by attention or arousal (Buhusi & Meck, 2005; Wearden, 2013). It could also be related to the putative positive valance of abstract symmetry (Makin, Pecchinenda, & Bertamini, 2012).

### An experiment on number of folds

Given that symmetry apparently elongates subject duration (at least in the sub-second range) we investigated whether this is further modulated by the number of axes of reflection (folds). Example stimuli with 1-5 folds are shown in Figure 1. These dot patterns have previously been used in Event Related Potential (ERP) research by Makin et al. (2016).

Such dot patterns are valuable because their *perceptual goodness* (i.e. salience or strength of the gestalt) can be quantified with *holographic weight of evidence model* (Nucci & Wagemans, 2007; van der Helm & Leeuwenberg, 1996). The holographic model assigns a ‘W-load’ to dot patterns. This is determined with the W = E/N formula, where W is perceptual goodness, E is the number of ‘holographic identities’ in a pattern, and N is the number of dots. A ‘holographic identity’ is a substructure with the same regularity as the whole (e.g. a reflected pair of dots in a global reflection). As explained by van der Helm (2011), there is then a non-monotonic relationship between W and the number of folds (1F W = 0.5, 2F W = 0.75, 3F W = 0.667, 4F W = 0.875, 5FW = 0.8). Makin et al. (2016) found that W predicted the amplitude of neural symmetry response successfully (the expected dip at 3 and 5F was evident in the at around 400 ms). We therefore predicted that the subjective elongation would also scale with W (as determined by the number of folds in our stimuli).

## Methods

### Participants

There were 40 participants (age = 18-55 years, 9 males, 1 left-handed). All had normal or corrected-to-normal vision. The experiment was approved by the Ethics Committee of the University of Liverpool and were conducted in accordance with the Declaration of Helsinki (2008).

### Power analysis

The smallest published effect of symmetry on subjective duration had an effect size of η_p_^2^ =0.194 (Palumbo, Ogden, et al., 2015). This was estimated from a sample of 24 participants. With this effect size, observed power was just 0.616. If this is the true population effect size, we have a high chance of obtaining a false negative with another sample of 24 (p = 0.384). However, with 40 participants, we had an 0.85 chance of replicating the basic effect of symmetry on subjective duration. The required sample size for finding a modulatory effect of folds is unclear because we do not have an a priori estimate of effect size for this.

### Apparatus and stimuli

Participants sat darkened cubicle 140 cm back from a 29 × 53 cm LCD monitor. The experiment stimulus was controlled using PsychoPy software (Peirce, 2007). Participants entered their responses on keyboard when prompted. A different pattern was generated on every trial, so no participant ever saw the same exemplar twice. The stimulus construction algorithm is described in detail Makin et al. (2016) and (Palumbo, Bertamini, & Makin, 2015). There were around 510-540 dots in each pattern. The mean number of dots varied slightly with the *N* axis. The SD of N dots also increased with *N* axis from 16 dots in the random conditions to around 50 in the 5-fold condition. Because the folds created some orientation cues, well included there were 1-5 folds in the random conditions (Figure 1 top row). The circular frame was approximately 5 degrees in diameter, as in Makin et al. (2016).

**Figure 1.**
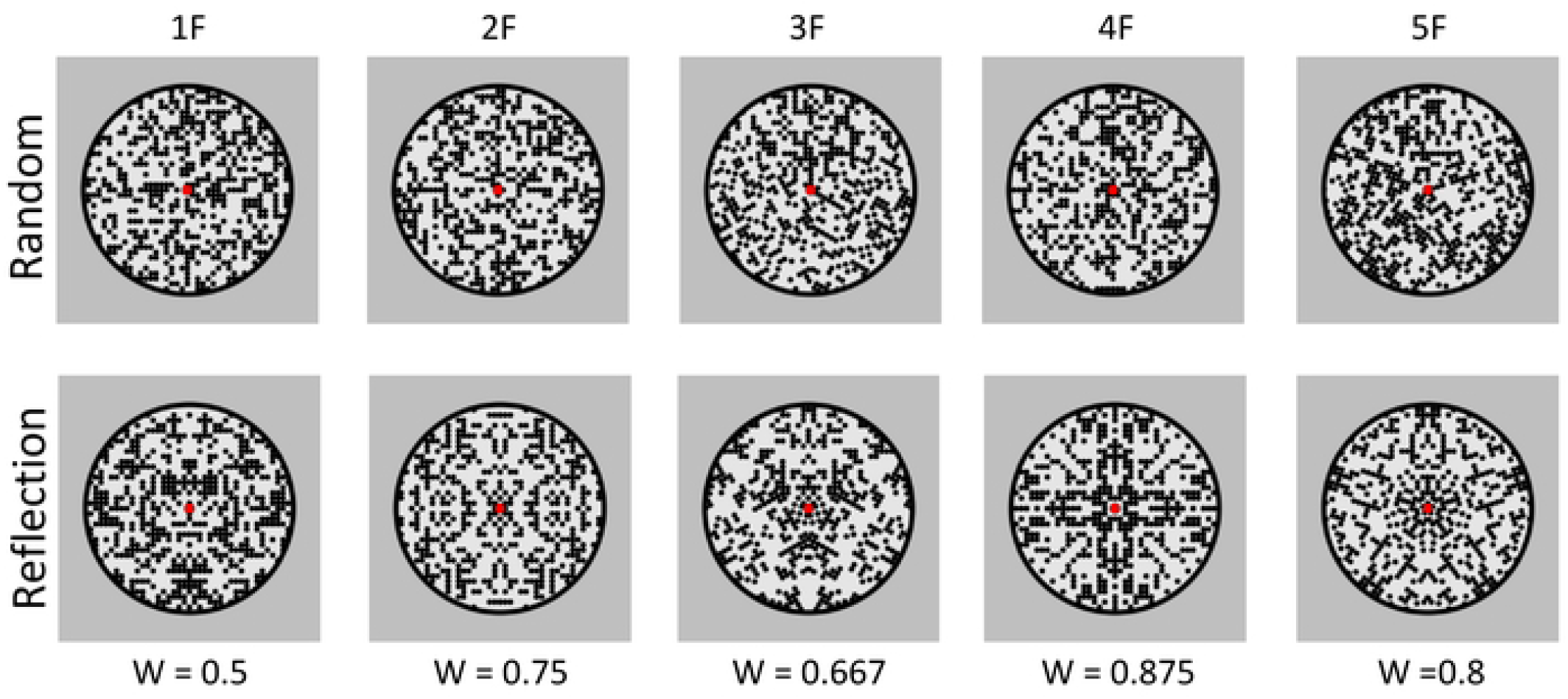
Example stimuli and W scores from the holographic model.

### Procedure

The experiment used a 2 x 5 x 3 within-subjects design. The factors were Regularity (reflection and random), Number of folds (1, 2, 3, 4 or 5) and Presentation duration (0.4, 0.6 and 0.8 seconds). There were 10 repeats of each *experimental trial* condition, and thus 300 experimental trials in total.

There were also an additional 300 *filler trials*. The filler trials were also fixed in terms of regularity (reflection, random) and folds (1-5). However filler trial duration was selected at random from 0.1 to 0.9 seconds. The filler trials prevented participants from overlearning the three durations of the experimental trials. At the end of each trial, participants used the mouse to rate duration estimates from 0 seconds to 1 second. The scale had 0.01 second increments, so we effectively continuous.

### Analysis

Most analysis was conducted on the Experimental trials. For each participant, the average duration estimate in was obtained in all 30 conditions. These were submitted to 2 Regularity X 3 Duration X 5 Folds repeated measures ANOVA. All 30 variables were normally distributed according to Shapiro-Wilk test (p > 0.07) and 29/30 were normally distributed according to Kolmogorov-Smirnov test (p > 0.2).

We also analysed the Filler trials. Here the actual duration varied randomly between 0.1 and 0.9 seconds (so there was no systematic Duration factor). These were submitted to 2 Regularity X 5 Folds repeated measures ANOVA. Again all 10 variables were normally distributed according to both Shapiro-Wilk (p > 0.614) and K-S tests (p > 0.097). The Greenhouse-Geisser correction factor was employed when the assumption of sphericity was violated (Mauchly’s test p < 0.05).

### Open science policy

Results are available on Open Science Framework, along with the PsychoPy code for running the experiment (https://osf.io/a3u6e/). Other researchers can use this material freely.

## Results

Duration estimates from the experimental trials are shown in Figure 2A. Although participants typically underestimated duration, subjective duration increased alongside actual duration (F (1.129, 44.021) = 217.481, p < 0.001, η_p_^2^ = 0.848). There was no effect of N axes (F < 1) and no effect of Folds (F < 1). There was no interaction between Regularity and Duration (F (2,78) = 1.835, p = 0.167) and no other interactions (F < 1).

The filler trials were less well controlled because actual duration varied randomly between 0.1 and 0.9 seconds. However, we can still assess effect of Regularity and Folds on mean duration estimates in the filler trials. As can be seen in Figure 2B, there were no effects of Regularity (F < 1) or Folds (F (4,156) = 1.210, p = 0.309) and no Regularity X Folds interaction (F (4,156) = 1.409, p = 0.233).

In the filler trials, most participants showed a strong correlation between actual duration and subjective duration. This shows that they were sensitive to filler trial duration. However, mean correlation coefficients was very similar whether patterns were symmetrical (Mean Pearson’s r = 0.623, SD = 0.164) or random (M = 0.624, SD = 0.613). This difference was not significant (t (39) = 0.115, p = 0.908). Therefore, there was no evidence that visual properties of the patterns altered performance.

**Figure 2.**
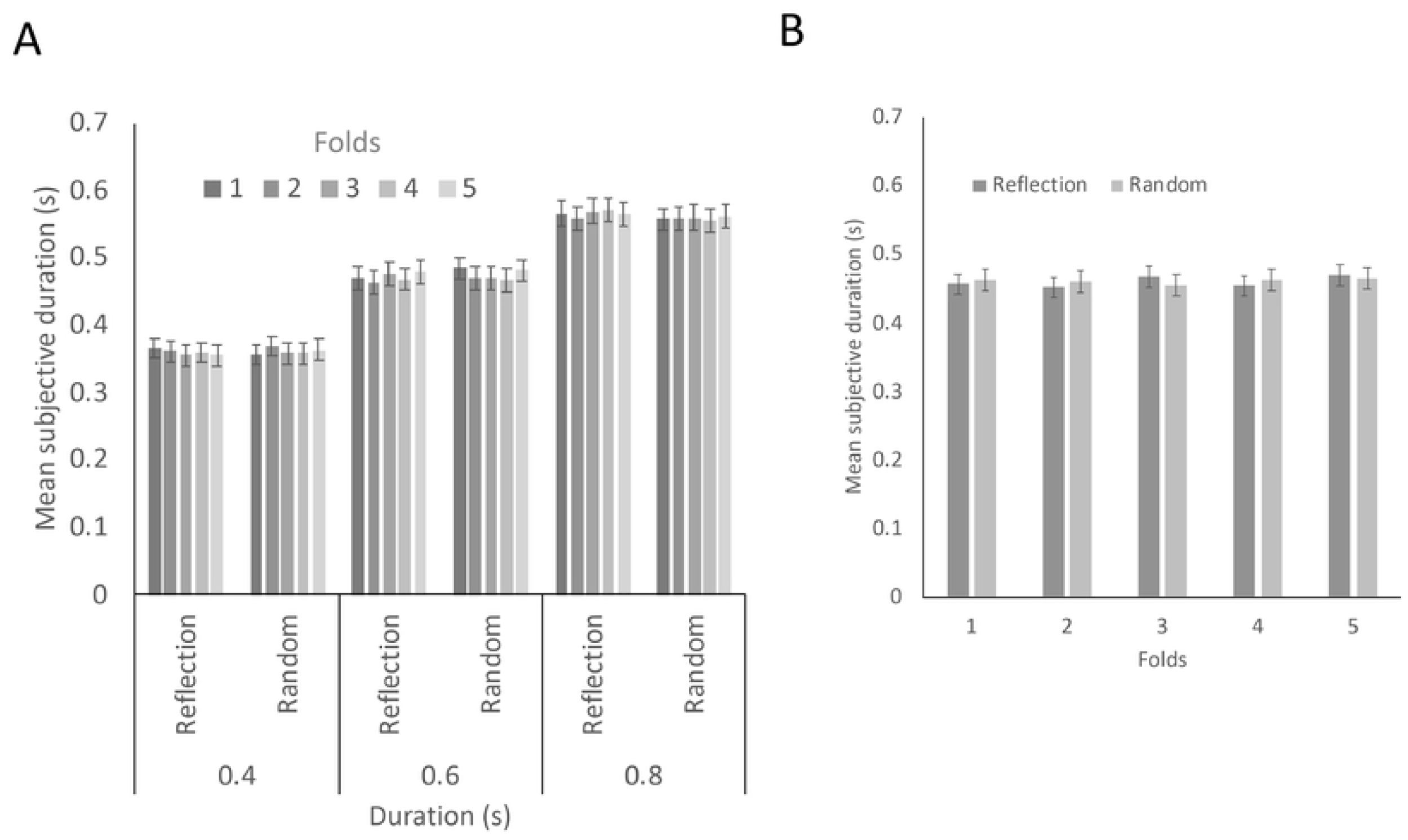
Results. A) Mean subjective duration in experimental trials as a function of actual duration. This is separated into reflection and random conditions with 1-5 folds. B) Mean subjective duration in filler trials in reflection and random conditions with 1-5 folds. Error bars = +/- 1 S.E.M.

## Discussion

Despite our predictions, there was no evidence that symmetrical dot patterns were judged longer than random dot patterns, and no evidence that this effect was further modulated by the number of folds. The basic effect of symmetry on subjective duration already reported by Palumbo et al. (2015), Ogden et al. (2016) and Sasaki and Yamada (2017) was not replicated. We are thus less confident that this is a robust effect. Arguably leaving it unpublished would be to contribute to the file draw problem, and all perversions of science that follow (Brysbaert, 2019; Laws, 2013).

Null results are often hard to interpret (hence they go unpublished). However, null results are scientifically relevant, especially when there is no obvious flaw in the study (such smaller sample sizes or very noisy data). For instance, null results from an underpowered experiment are less interesting. However, our experiment was not statistically underpowered by conventional standards (a priori power > 0.85), and there was no hint of a trend in the expected direction.

A failed replication in experimental psychology could also be scientifically uninteresting if the stimuli differ from original experiments in some obvious way that (with hindsight) was always very likely abolish the expected effect. For instance null result might be uninteresting if stimulus contrast was much reduced, and the visual effect under investigation is known to require high contrast stimuli. However, this does not apply here. Our dot patterns differed from the checkerboards in Palumbo, Ogden et al. (2015) and Ogden et al. (2016), and from the dot lattices in Sasaki and Yamada (2017). However, the symmetry was *more* salient here, and if anything, the effect of symmetry on duration should have been larger (Makin et al., 2016; van der Helm & Leeuwenberg, 1996). Furthermore, there is no reason to believe our dot patterns were more/less arousing, distracting or aesthetically pleasing than those used previously.

A failed replication could also be scientifically uninteresting if the task had changed in an obviously problematic way. Palumbo, Ogden et al. (2015) and Ogden et al. (2016) used a verbal estimation task, where participants entered judgements using the keyboard. Instead, our participants entered judgments using the mouse to scroll a cursor left and right along a continuous scale. This may have resulted in less ‘quantizing’, where participants tend to give round number estimates (such as “500 ms” or “1000 ms” Wearden, 2015). This extra precision of our response scale could have introduced ceiling effects and masked the effect of symmetry. However, the effect of actual duration on subjective duration was comparable to previous work (Current study η_p_^2^ = 0.85, Ogden et al. 2016, η_p_^2^ = 0.88, Palumbo, Ogden et al. 2015, η_p_^2^ = 0.76). This suggests that suppressive ceiling effects were not enhanced the current study.

In summary, this is a pure failure to replicate, without clear explanation. We believe such failures to replicate should not end up in the file drawer. Timing researchers should not have the mistaken impression that the effect of symmetry on subjective duration has been established without exception.

## Acknowledgements

This project was part funded by an ESRC grant award to Alexis Makin (ES/S014691/1)

